# Explainable multi-view framework for dissecting intercellular signaling from highly multiplexed spatial data

**DOI:** 10.1101/2020.05.08.084145

**Authors:** Jovan Tanevski, Ricardo Omar Ramirez Flores, Attila Gabor, Denis Schapiro, Julio Saez-Rodriguez

## Abstract

The advancement of technologies to measure highly multiplexed spatial data requires the development of scalable methods that can leverage the spatial information. We present MISTy, a flexible, scalable and explainable machine learning framework for extracting interactions from any spatial omics data. MISTy builds multiple views focusing on different spatial or functional contexts to dissect different effects, such as those from direct neighbours versus those from distant cells. MISTy can be applied to different spatially resolved omics data with dozens to thousands of markers, without the need to perform cell-type annotation. We evaluate the performance of MISTy on an *in silico* dataset and demonstrate its applicability on three breast cancer datasets, two measured by imaging mass cytometry and one by Visium spatial transcriptomics. We show how we can estimate interactions coming from different spatial contexts that we can relate to tumor progression and clinical features. Our analysis also reveals that the estimated interactions in triple negative breast cancer are associated with clinical outcomes which could improve patient stratification. Finally, we demonstrate the flexibility of MISTy to integrate different kinds of views by modeling activities of pathways estimated from gene expression in a spatial context to analyse intercellular signaling.

## Introduction

Highly multiplexed, spatially resolved data is becoming available at an increasing pace thanks to recent and ongoing technical developments. In contrast to dissociated single-cell data, this data informs us on the cell-to-cell heterogeneity in tissue slices while conserving the arrangement of cells^1^. Therefore, each cell can be studied in its microenvironment. We can observe the spatial distribution of the expression of markers of interest, their interactions within the local cellular niche and at the level of tissue structure. All these aspects provide an excellent platform to gain better insight into multi-cellular processes, in particular cell-cell communication.

The proliferation of spatial technologies leads to the generation of large amounts of data. Different technologies allow for measuring different types of molecules with varying resolution, capturing different areas of tissue with diverse numbers of readouts. Immunofluorescence-based methods allow detection of the expression of tens to hundreds of proteins at subcellular resolution^2–4^ and hundreds to potentially thousands of RNA species at single-cell resolution^5^. Mass spectrometry-assisted methods enable detection of the expression of a high number of proteins at the resolution of tissue patches^6,7^ and tens of markers at subcellular resolution^8,9^, and over hundred metabolites at cellular and subcellular resolutions^10,11^. Finally, barcoding-based approaches^12^, facilitate the measurement of genome-wide expression at a resolution of hundreds of microns, i.e., several cells, and are being further developed to increase the resolution to below ten microns^13,14^. Complementally, we are also witnessing the rapid development of methods for spatial localization that combine limited amounts of spatially resolved data with richer, but dissociated single-cell data^15–19^, which can alleviate the various shortcomings of the technologies. Therefore, there is a need for methods to analyse large amounts of rich and spatially-resolved data in order to discover patterns of expression, interaction and cell functions. In fact, this has been identified as one of the grand challenges in single-cell data science^20^. These methods should ideally be able to handle the variety of produced data and scale well with future technology improvements.^21–24^

Currently, there is a limited number of computational methods available for the analysis of high-resolution spatially-resolved data^25^. One group of methods focuses on the analysis of the significant patterns and the variability of expression of individual markers^21–24^ to describe the landscape of expression within a tissue. Another group of methods considers, more broadly, the analysis of the interactions between the markers within different spatial contexts, that is the expression in the directly neighboring cells or the effect of the expression of a marker in the broader tissue structure. The methods within the latter group focus mainly on identifying interactions in the local cellular niche, by establishing the statistical significance of the distribution of automatically identified cell types in the neighborhood of each cell^26–31^. These methods assume a fixed form of nonlinear relationship between markers or have a predefined set of spatial contexts which can be explored. Spatial Variance Component Analysis (SVCA)^32^, for example, goes a step further by examining intercellular interactions by decomposing the source of variation to three fixed spatial contexts: intrinsic, environmental and intercellular effects.

We introduce here a Multiview Intercellular SpaTial modeling framework (MISTy), an explainable machine learning framework for knowledge extraction and analysis of highly multiplexed, spatially resolved data. MISTy facilitates an in-depth understanding of marker interactions by profiling the intra- and intercellular relationships. MISTy is a flexible framework able to build models to describe the different spatial contexts, that is, the types of relationship among the observed expressions of the markers, such as intracellular regulation or paracrine regulation. For each of these contexts MISTy builds a component in the model, called a view. MISTy allows for a hypothesis-driven and flexible definition and composition of views that fit the application of interest. The views can also capture functional relationships, such as pathway activities and crosstalk, cell-type specific relationships, or focus on relations between different anatomical regions. Each MISTy view is considered as a potential source of variability in the measured marker expressions. Each view is then analyzed for its contribution to the total expression of each marker. The measured contribution points to the relevance of a potential source of interactions coming from the different spatial contexts and is estimated from the view specific models. Our approach is modular, easily parallelizable and thus scalable to samples with millions of cells and thousands of measured markers. While inspired by other approaches^21–24,26–28^ to explicitly model the spatial component of the data, MISTy’s approach is unique: First, it models the complete measured expression profile and interactions instead of analysing spatial patterns of single markers. Second, it is not limited to fixed predefined sources of variation, aggregation or representation of the data, but allows for the flexible construction of models to analyse spatial data. Third, it does not require to annotate the cell-type, state, or any other feature of the spatial unit (cell or spot).

We validated MISTy on *in silico* data generated by a custom algorithm. We further applied our framework on two different Imaging Mass Cytometry (IMC) datasets consisting of 46 and 720 breast cancer biopsies respectively. On these data sets, we demonstrated how MISTy outperforms available methods by recapitulating previous results and at the same time adding interpretation and new insights. This enabled us to discover intra- and intercellular features in triple negative breast cancer that are associated with clinical outcomes. To our knowledge, this is the first method available to connect spatially resolved single cell measurements to clinical outcome without the use of cell type annotation. Finally, MISTy can extract knowledge about the interactions among signaling pathways and ligands expressed in the microenvironment from different spatial views. We demonstrate this on spatial transcriptomics data of breast cancer. These case studies illustrate the flexibility of MISTy as a framework to define exploratory and hypothesis-driven workflows for the analysis of diverse types of spatial omics data in basic and translational research.

## Results

### MISTy: Multiview intercellular spatial modeling framework

MISTy is a late fusion multiview framework for the construction of a domain-specific, explainable model of the expression of markers (Figure 1). For each marker of interest in a sample, we can model cell-cell interactions coming from different spatial contexts as different views. For example, the first and main view, containing all markers of interest, is the intraview, where we relate the expression of other markers to a specific marker of interest within the same location. To capture the local cellular niche, we can create a view that relates the expression from the immediate neighborhood of a cell to the observed expression within that cell; we call this view a juxtaview. To capture the effect of the tissue structure, we can create a view that relates the expression of markers measured in cells within a radius around a given cell, and we call this view a paraview (see Methods). Importantly, MISTy is not limited to the abovementioned views. Other views can be added to the workflow that can offer an insight about relations coming not only as a function of space. For example, views can focus on interactions between different cell types, interactions within specific regions of interest within a sample or a higher level functional organization.

**Figure 1.**
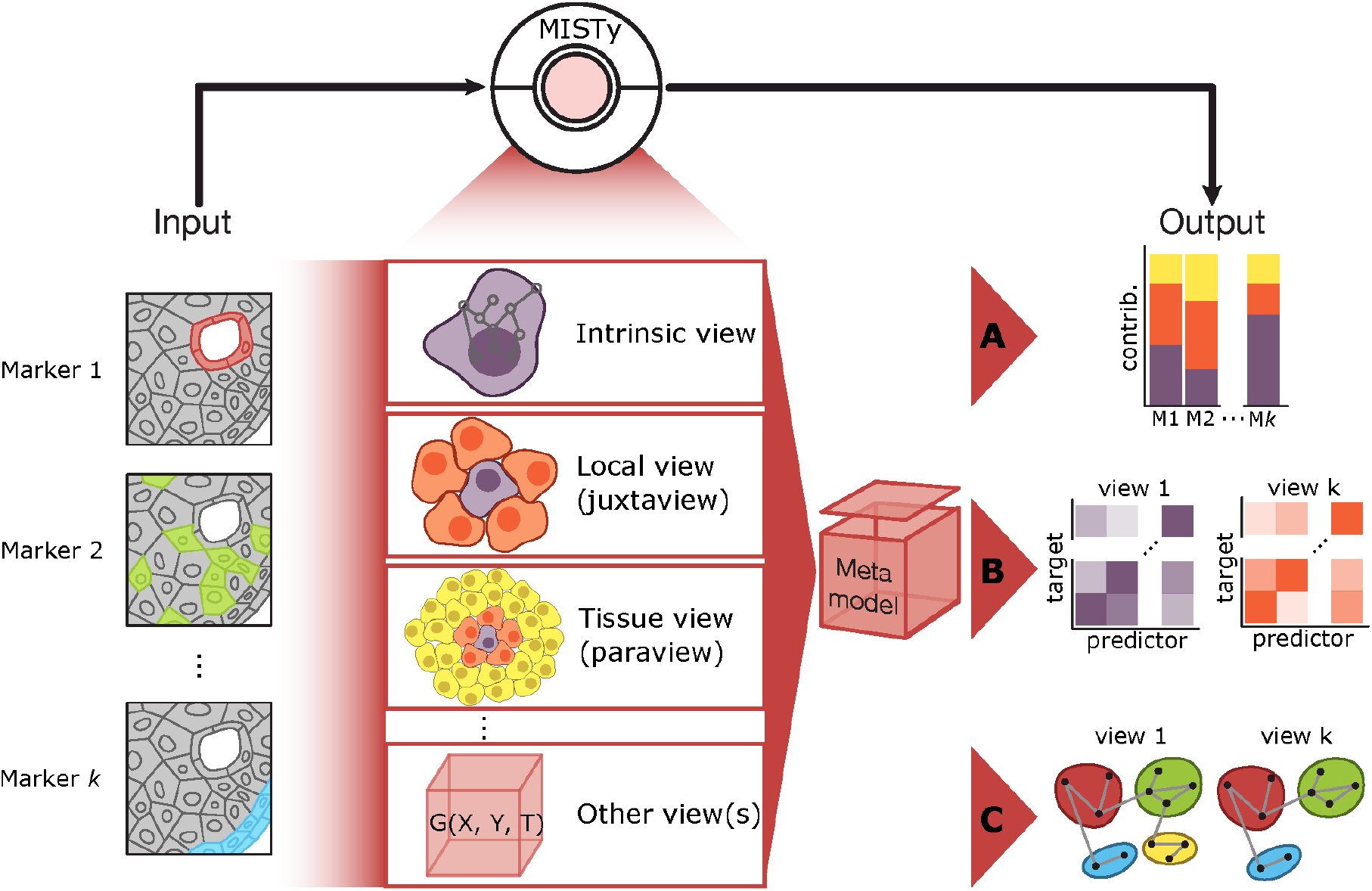
MISTy: An explainable multi-view framework for modeling intercellular interactions from highly multiplexed spatial data. MISTy models marker relations coming from different spatial views: intrinsic (intraview), local niche view (juxtaview), the broader, tissue view (paraview), or others, based directly on marker expressions or derived typology or functional characterisations of the data. At output, (A) MISTy extracts information about the contribution of different spatial views to the expression of markers in each cell. (B) MISTy also estimates the markers’ interactions coming from each view that explain those contributions. (C) These results can be described qualitatively as communities of interacting markers for each view.

Formally, we consider a matrix [*Y*]_*u,i*_ where each column represents a marker (*i* = 1 .. *n*) and each row is a spatial location (*u* = 1 .. *L*). *Y*_*.,i*_ is the vector made by all observations of the marker *i*. MISTy models its expression as

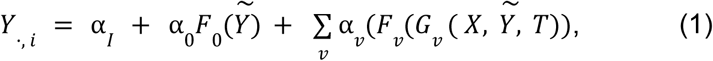

where 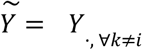, i.e. all markers except the target marker. *F*_*v*_ are models constructed by a machine learning algorithm (in this work we consider *F* to be Random Forests^33^) for each view *v*. *G* are domain-specific functions that transform the data to generate informative variables (features) from the expression *Y* at the corresponding spatial localization *X*. Optionally, *G* can depend on other specific properties *T*, such as prior-knowledge expressed as annotated functions, regions or cell-types. The *G* functions can be used to generate alternative views that can be inputs to the model function *F*. For example, given gene expression data, a function *G* can be used to infer pathway activities at each location. The corresponding variables can be input to MISTy to relate the activities of pathways at each location with those from a broader spatial context. Finally, α are the late fusion parameters of the meta-model that balances the contribution of each view to the prediction.

The above model is trained in two steps. First, the models for each view are trained independently. Second, we estimate α parameters of the meta-model after training the view-specific models independently, by linear regression.

MISTy always models a fixed, intraview 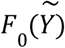 as a baseline view that is independent from the spatial localization of the cell. It is biologically expected that this intraview will be able to capture most of the variance of the expression of the measurements: the effects on the measured markers from outside of the cell are normally lower than the effects of the interactions and regulation coming from within the cell itself^34^. By design, our focus is to distinguish the non-intrinsic effects from the intrinsic baseline and estimate important interactions that supplement the explanation of the overall expression. To this end, other intercellular views are then added to 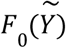. The user can add a number *v* of additional, intercellular views, and separate the effect of each view for each marker on the improvement in predictive performance of the multiview model. We use the improvement in predictive performance of the models as a proxy to estimate their potential as sources of interactions that can be further explored by extracting feature importances, as outlined in the following. The contribution of each view is captured by the late fusion parameters α of the meta-model.

MISTy is a general framework and can construct models for the functions *F* with any algorithm that fulfills two requirements. First, the algorithm should construct ensemble models, with constituents trained on a bootstrap sample (bag) from the data. Second, they should be or consist of explainable models. The first criteria guarantees the unbiased use of the measurements in both steps of model training. The predictions of the constituents of an ensemble model can be made on portions of the data (out-of-bag) that were not used for their training. The second criteria means that a global explanation of the model or the importances of the features can be obtained post-hoc from the trained models. As proof of principle, in this work we use Random Forest^33^ for *F*. Random forests are well known, robust and flexible models fitting the two criteria outlined above and have been shown to achieve good performance in various application areas. The feature importances for Random Forest models can be explained by the reduction of variance in the constituent trees.

At the first level, the meta-model can be interpreted in two different ways. First, to answer the question of how much the intercellular views improve the prediction of the expression in addition to the intracellular view. This can be achieved by comparing the predictive performance of a single intracellular view vs all views combined in a meta-model. Second, by comparing the values of the fusion parameters, we can investigate how much the individual views contribute to explaining the marker expression that led to the aforementioned improvement in predictive performance (Figure 1A).

At a second level, given this information, we can further analyse the feature importances. For each target marker, we can inspect each view-specific model and analyse how important is the contribution of each marker in that view to the prediction of the expression of the target marker (Figure 1B). Thus, we estimate the interactions among the markers from the individual marker and view specific models. However, for every marker, the statistical significance of the contribution of the view-specific models in the meta-model is explicitly taken into account when calculating the importances (see Methods, Importance weighting and result aggregation). These importances correspond to potential relations between the predictor and the target marker in the specific spatial or other context modelled by the corresponding view. MISTy outputs the estimated importances of significant marker relations. Since these relations are based on the importance of a marker in predicting the target they cannot be assumed to be directly causal nor directional. The relations between markers may occur through a network of intermediate interactions in the specific biological context, which can be further explored by enrichment of these relations using curated databases of intra- and intercellular interactions (Figure 1C).

Finally, if multiple samples are available during the analysis, the relations from individual samples are aggregated to produce robust results (see Methods). By aggregation, we accentuate consistently inferred interactions from individual samples and reduce the number of false positive interactions.

### *In silico* performance

We first assessed the performance of MISTy to reconstruct *in silico* intra- and intercellular interaction networks. For this, we created a tissue simulator that can mimic the interactions of different cell-types through ligand binding and subsequent signaling events (Figure 2A; see Methods). The dynamic model simulates the production, diffusion, degradation and interactions of 11 molecular species including ligands and intracellular signaling proteins (see Methods) until a steady state distribution of markers are reached.

**Figure 2.**
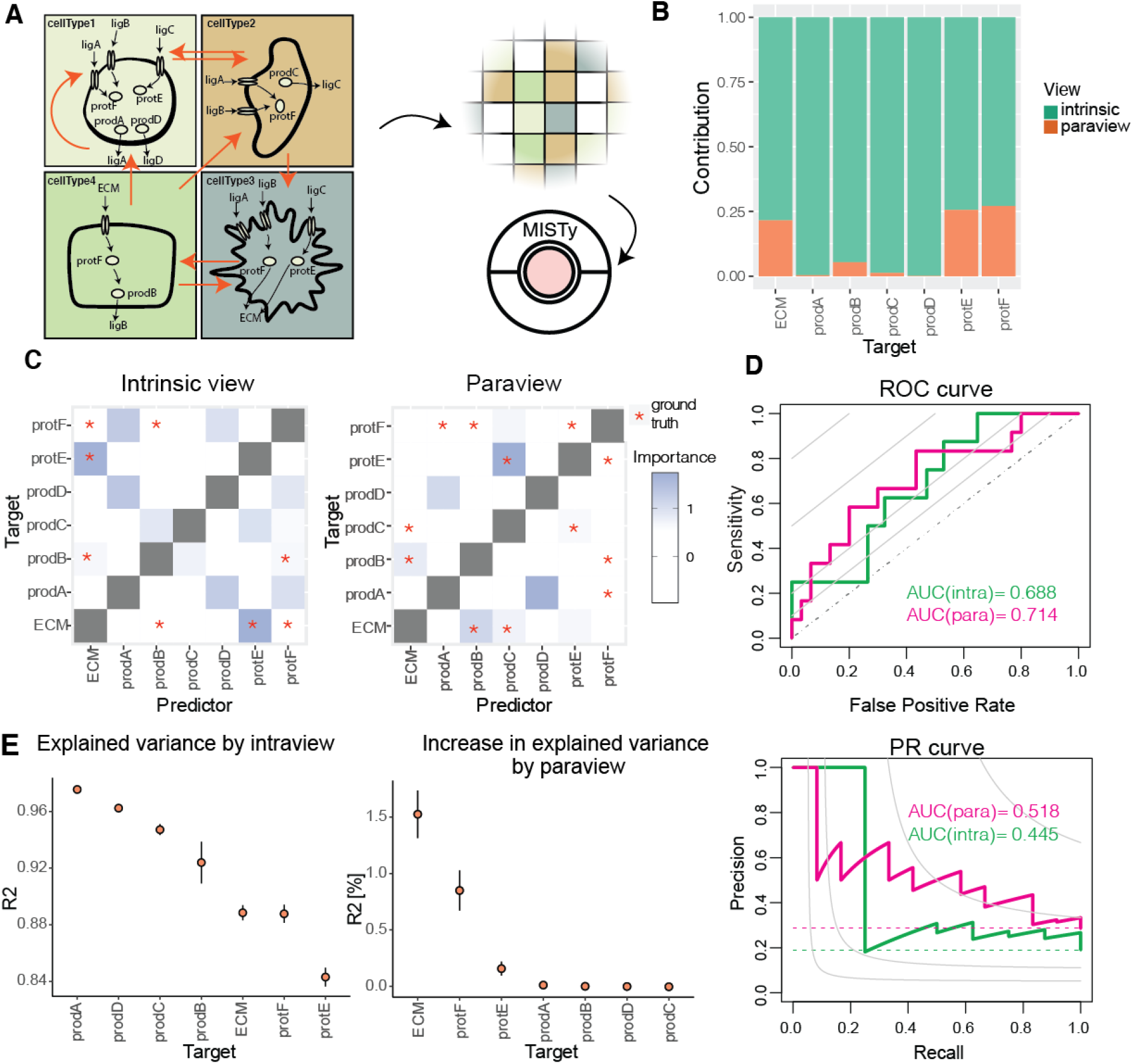
Evaluating MISTy on in silico data. (A) MISTy was evaluated on the task of reconstruction of simulated interaction networks. Models of intra-(black arrows) and intercellular (orange arrows) interactions of four different cell types, arranged on a grid representing a tissue, were used to simulate measurements of 11 molecular species. (B) Contribution of each view to the prediction of the marker expressions in the meta-model. The stacked barplot represents normalized values of the fusion coefficients of the respective views for each marker. (C) Marker interactions in the intrinsic and paraview. Heatplot shows the interactions found by MISTy, red stars highlight the ground truth interactions. (D) Receiver operating characteristic (ROC) and precision-recall (PR) curves depicting the aggregate performance of MISTy on all ten samples for the intraview and paraview. The dashed lines represent the expected performance of an uninformed classifier, the gray iso-lines represent points in ROC space with informedness (Youden’s J statistic) equal to 0.1, 0.2, 0.5 and 0.8 and points in PR space with F1 measure equal to 0.1, 0.2, 0.5 and 0.8. (E) Explained variance of each marker by the intraview alone and the increase in explained variance by adding the paraview contribution.

The simulated values for every molecular species, at every location is recorded and passed as input to MISTy (Sup.Figure 1). Information about the different cell-types, their intracellular wiring or which cell type expresses which ligand is not given as input to MISTy. We use this procedure to estimate the robustness of MISTy to infer interactions.

The MISTy workflow for *in silico* data consists of two views, intracellular view and broader tissue structure view (paraview). When we compare the predictive performance of this model to a model with a single intraview, we see the highest improvement in predictive performance for the expression of markers ECM, prodE, prodF and prodB. The improvement is reflected in the observed contributions of the tissue structure view (Figure 2B). Markers that were not affected by environmental interactions prodA, prodD and prodC showed, as expected, no improvement in the paraview.

We evaluated the performance of MISTy to recover interactions among markers. MISTy identified strong importance between ECM and protE in the intraview and between prodC and protE in the paraview (Figure 2C). These two steps are part of the ECM production pathway involving cellType2 and cellType3. In the dynamical model, prodC produces ligC in cellType2, which diffuses on the lattice and activates protE in cellType3. In turn, protE activates ECM production in cellType3. We also see two interactions with high importance scores in the intraview: prodA - protF and prodA - protD. The former corresponds to the autocrine signaling in cellType1, where prodA produces ligA that activates protF. The latter interaction (prodA - prodD) shows that the co-expression of two markers can be identified as interaction.

The paraview captures three interactions: prodC - protE and prodC - ECM, which are participating in the cell-cell communication between cellType2 and cellType3 through ligC; and ECM - prodB corresponds to the process where ECM activates prodB through protF in cellType4. Although ProtF has a large paraview contribution (Figure 2B), we cannot see an important interaction that is responsible for that (Figure 2C), likely because protF is involved in various interactions across all cellTypes (protF is activated by ECM in CellType4, activated by ligB in CellType3 and by both ligA and ligB in CellType1 and 2).

Overall, the intraview alone explains a large amount of variance of the nodes that appears only in the intracellular space and do not diffuse (Figure 2E) and the paraview module increases the model accuracy for nodes that are activated by diffusing molecules in the intercellular space.

Across the individual samples, we observed variance in MISTy’s performance, in both the area under the receiver operating characteristic curve (AUROC) and the area under the precision-recall curve (AUPRC) (Sup.Figure 2). We aggregated the results from all layouts (see Methods section for details) and calculated the performance (Figure 2D). The aggregation strategy maximized the extracted knowledge available in the samples (AUROC= 0.688 and 0.714 and AUPRC = 0.445 and 0.518 for the intrinsic and the paraview, respectively). In summary, MISTy is able to reliably extract interactions in the *in silico* case study.

### Application to Imaging Mass Cytometry breast cancer datasets

#### Analyzing the importance of the tissue structure

As a first real-data case study, we applied MISTy to an Imaging Mass Cytometry dataset consisting of 46 samples of breast cancer across three tumor grades coming from 26 patients, with measurements of 26 protein markers^28^. We processed each sample independently, with 944 cells on average per sample, or 43434 single cells in total. We designed the exploratory MISTy workflow for this task to include three different views capturing different spatial contexts and providing a foundation for comparison with SVCA: An intraview, a view focusing on the local celular niche (juxtaview), and a view capturing the effect of the broader tissue structure (paraview), as detailed in the Methods section.

We aggregated the MISTy results from all samples and we found that the multiview model resulted in significant improvements in absolute value of variance explained of up to 23.6% over using the intraview alone, which accounted for an average of 23.9% of overall variance explained across all markers (Sup.Figure 3A). This is consistent with results obtained with SVCA, on the same data^32^. Highest improvement was detected for the markers pS6 (5.13% ± 4.99), CREB (4.10% ± 3.31) and SMA (4.08% ± 3.27) (Figure 3A). This is expected, since these three markers have distinct spatial distributions: pS6 represents “active” stroma present in distinct regions of the tumor microenvironment, SMA represents smooth muscle Actin, which is expressed in ductal structures and blood vessels; and CREB is a transcription factor commonly overexpressed and activated in tumor regions. The highest change in variance explained (23.6%) in a single sample was observed for CAIX, a marker of hypoxia. All top ranked markers by improvement found by MISTy are consistent with the highest improvement due to environmental effect in the results of SVCA.

**Figure 3.**
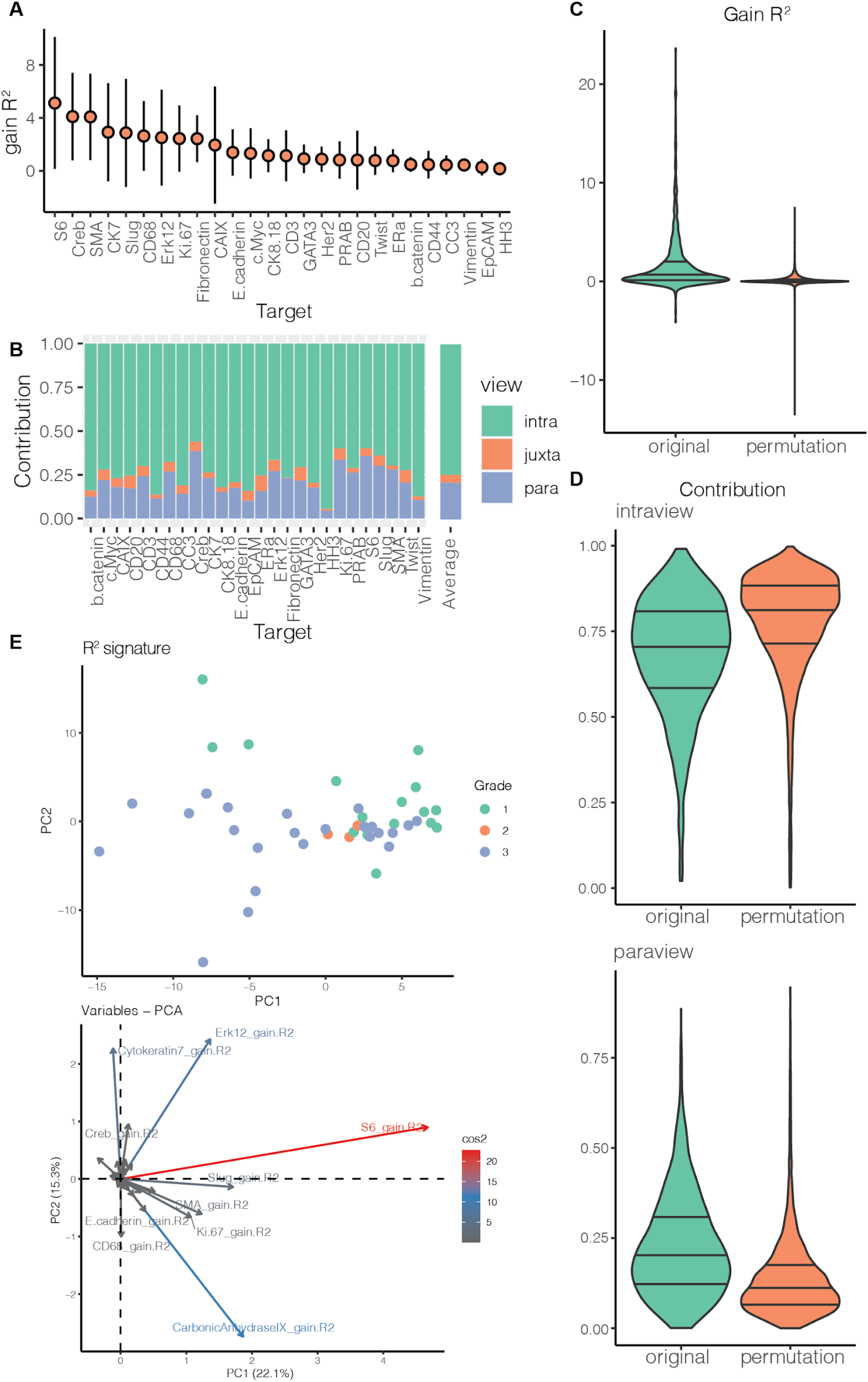
R^2^ signature and permutation analysis of IMC data from 46 breast cancer samples. (A) Improvement in the predictive performance (variance explained) for all samples when considering multiple views in contrast to a single, intraview (in absolute percentage points). (B) Relative contribution of each view to the prediction of the expression of the markers. (C) Distribution of improvement in variance explained when considering multiple views in contrast to a single intraview across all markers and samples with original cell locations and 10 random permutations. (D) Distribution of the relative contribution of the interview and the paraview to the prediction of the markers across all markers and samples with original cell locations and 10 random permutations. (E) First two principal components of the R^2^ signature of the samples and the importance of the variables of the signature in the principal component analysis.

We next analyzed the contribution of each view to the prediction of the multi-view model (Figure 3B). With MISTy, unlike SVCA, we were able to dissect the effect of the juxtaview and paraview. We find that a significant contribution (higher value of the fusion parameter in the meta-model) comes from the paraview compared to the juxtaview. This suggests a stronger effect from the broader tissue structure than from the immediate neighbours. The mean fraction of contribution to the prediction of the intraview was 74.5%, of the juxtaview 4.5% and of the paraview 21%. These results illustrate how MISTy can recapitulate previous findings without the need of single cell clustering and cell type annotation using prior knowledge^28^.

To investigate the importance of tissue structure for the modeling of spatially resolved single cell data, we performed a spatial permutation based analysis and compared the results obtained by MISTy. The coordinates of each cell in each sample were permuted 10 times. Subsequently, we ran the aforementioned MISTy workflow on the resulting 10 new datasets. The mean gain in variance explained for the permuted data across all samples and markers was 0.1% with 40% of values of the gain of variance explained less or equal to 0 compared to 1.7%, with only 15% of values of the gain of variance explained less or equal to 0 for the original data (Figure 3C), i.e. the availability of true tissue structure improves the performance of the mode significantly (*log*_10_*p* ≪ − 10; one-sided Wilcoxon rank-sum test of the distributions of the gain in variance explained between original and perturbed data)). The estimated contribution of the juxtaview and paraview for the permuted datasets was much lower than for the original dataset (*log*_10_*p* = − 2. 45and *log*_10_*p* ≪ − 10 respectively; one-sided Wilcoxon rank-sum test), and often nearly absent. In addition there were significantly higher contributions of the intraview than for the original dataset (Figure 3D, Sup.Figure 3B,C). For the permuted dataset the mean baseline variance explained over all samples and markers by using only the intraview was found to be consistent, i.e. remained the same as for the original dataset (23.9%).

Subsequently, we analyzed our results by the spatial variance signature (R^2^ signature) of each sample. We defined the R^2^ signature of the MISTy results for each sample by concatenating the estimated values of the variance explained using only the intraview, the variance explained by the multiview model and the gain in variance explained for each marker. The signature vector for each sample is therefore 78 (26 markers × 3 views) dimensional, and allows to compare the results of MISTy to SVCA. Using the first two components of the principal component analysis (PCA) of the R^2^ signature, we identified a structure in the samples that is driven by the tumor grade, which is consistent with the findings of SVCA (Figure 3E). The two first principal components of the R^2^ signatures of MISTy captured 37.4% of the variance of the samples compared to 30% with the spatial variance signature of SVCA. Inspecting the importance of the R^2^ signature components for the PCA analysis (Figure 3E), we observed that the structure of the results can be explained by the gain in variance explained, which points again to the relevance of the spatial component of the data. In particular the gain for markers pS6, CAIX, Erk12 and Cytokeratin 7 were found to be the highest, suggesting that proliferation, hypoxia and loss of structure in different grades of tumors is significantly affected by change in regulation as a result of intercellular interactions. Collectively, these results support the importance of the tissue structure for the expression of proteins at the single cell level.

While the R^2^ signature allows us to analyze the differences between the samples based on predictive performance only, more mechanistic insights can be obtained by a more detailed signature at the level of the estimated importances. The importance signature is generated by concatenating the estimated importances for each predictor-target marker pair from all views. The aggregated importances are weighted by the estimated relevance of the results (see Methods). In this case we created a 26 marker × 26 marker vector for 3 views (2028 dimensions). The signature vector for each sample is therefore larger but more informative and focused on interactions. The structure in the results, driven by the tumor grade, can also be observed when visualizing the first two principal components of the importance signature (Figure 4A). Due to richer information they account for less (14.5%) of the variance of the samples compared to the R^2^ signature. By inspection of the importance of the signature components, we observed that in the two first principal components the most significant interactions that can account for the observed structure and differences among the samples come from the broader tissue view.

**Figure 4.**
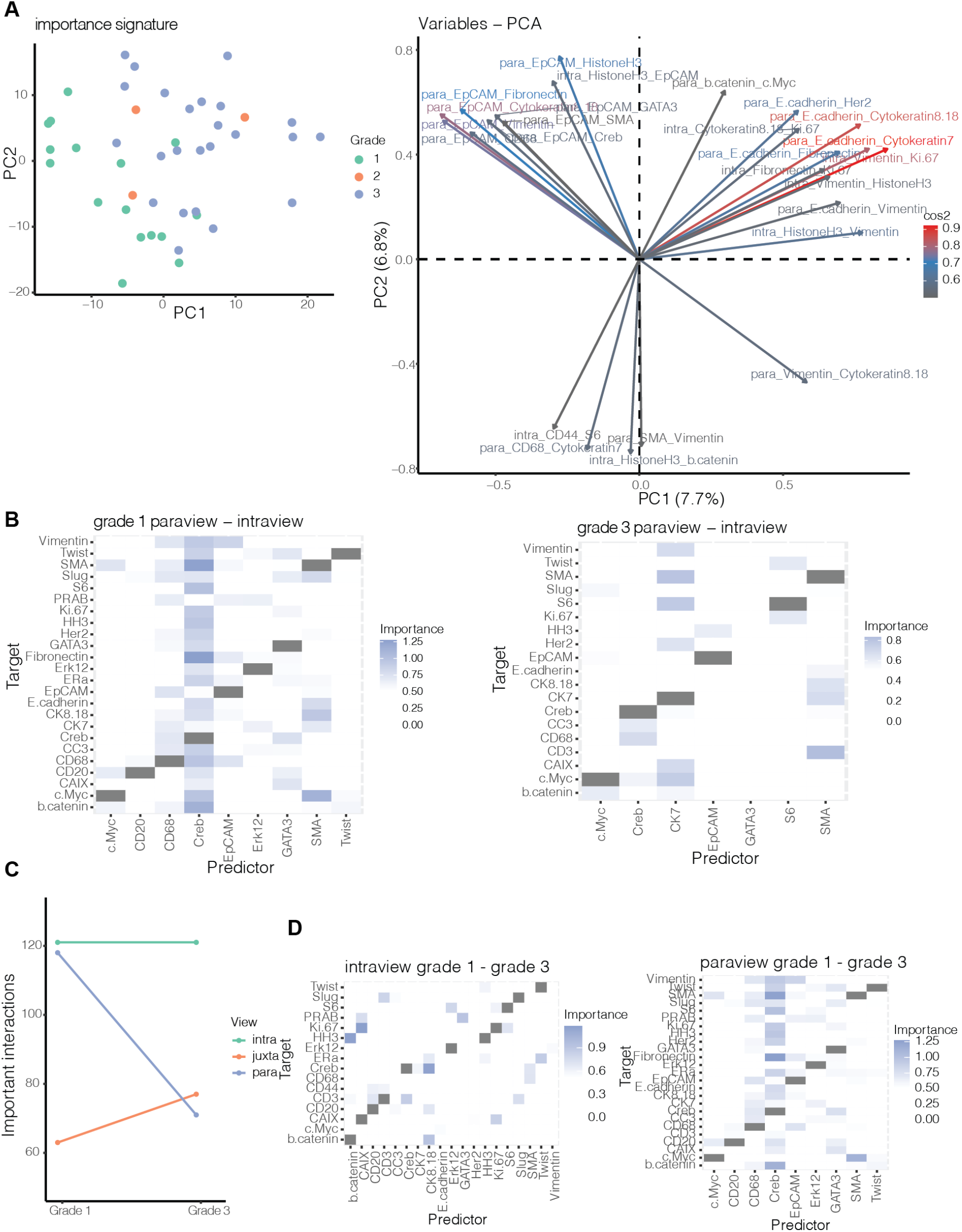
Importance signature and contrasts of IMC data from 46 breast cancer samples. (A) First two principal components of the importance signature of the samples and importance of the variables of the signature in the principal component analysis (25 variables with the highest square cosine shown). (B) Intragroup contrast of importances of marker expression as predictors of the expression of each target marker between the intraview and paraview for grade 1 samples and between the intraview and paraview for grade 3 samples. (C) Change of total number of estimated important interactions per grade (Importance ≥ 0.5) (D) Intergroup contrast of importances of marker expression as predictors of the expression of each target maker for the interview and for the paraview between grade 1 and grade 3 samples.

To confirm that the structure of the samples can be observed complementary as the result of accounting for the spatial component of the data and is not simply a result of the intrinsic expression of the markers, we performed PCA on the samples as represented by the mean expression of the markers across all cells. While the separation of the samples by grade is observable when visualizing the first two principal components (accounting for 63.9% of the sample variance), the importance of the markers that account for this separation are more uniform and different from the components of the R^2^ signature (Sup.Figure 3D).

#### Stratifying samples and highlighting intergroup differences

By grouping the samples by grade, we further analysed the robust intercellular features of tumor samples. Since only a small number of samples came from grade 2 tumors, we considered only grade 1 and grade 3 tumor samples. In grade 1 samples we observed the highest gain of variance explained for markers Cytokeratin 7 (5.6% ± 4.84), SMA (4.14% ± 3.35) and CREB (3.8% ± 2.57). In grade 3 samples we observed the highest gain of variance explained for pS6 (6.95% ± 5.09), SMA(4.43% ± 3.39), and CREB (4.26% ± 3.52). While the importance of Cytokeratin 7 and pS6 for separation of the sample by grade was already apparent from gain in variance explained (Figure 3E), with the importance signature we can also highlight the contributing interactions.

We further compared the aggregated results by contrasting the important interactions between the same views intragroup and intergroup. Due to the higher overall contribution of the paraview compared to the juxtaview (Figure 3B), we analyzed the important interactions that can be extracted from the paraview model that have not been found as significant for the intraview model, i.e. capture interactions coming only from the context of the broader tissue structure. In grade 1 samples we observed predictor markers with high importance for many target markers: the transcription factor markers CREB and GATA3, the immune cell marker CD68 and the myoepithelial marker SMA. In grade 3 samples we observed a pronounced decline of the number of important interactions coming from the paraview compared to grade 1 samples, with most important predictor markers being Cytokeratin 7 and SMA (Figure 4B). This is likely due to loss of normal tissue architecture in grade 3 samples. In other words, normal tissues are highly structured, and the underlying tissue structure is critical to perform tissue relevant functions. Advanced tumors create tissue structures that are dominated by tumor cells and thus are not as dependent on cellular crosstalk and organization.

The loss of signaling during tumor progression is apparent when comparing the results view by view for the different tumor grades. The intergroup and view focused contrasts outline the interactions that were estimated as important in grade 1 samples and not important in grade 3 samples. While we observed a loss of number of important interactions from the intraview, the loss of important interactions from the paraview is higher (Figure 4C, D).

#### Linking estimated interactions to clinical features

To highlight the ability to associate MISTy results with clinically-relevant features, we analyzed a breast cancer imaging mass cytometry dataset with outcome data, based on 415 samples from 352 patients (see Methods section for sample selection)^35^. As with the previous dataset, we processed each sample with MISTy independently and used the exploratory MISTy workflow with three views capturing different spatial contexts: intraview, juxtaview, and paraview.

The amount of important interactions, as with the previous breast cancer data set, decreased with tumor progression based on grading (Figure 5A) across all three views. The highest median improvements were detected for the same markers as shown in the previously described breast cancer data set showing reproducibility across multiple sample cohorts (Sup.Figure 4). Visualization of the network communities based on the estimated importances of the predictor - target pairs from the juxtaview for grade 1, 2 and 3, highlights the rewiring of the tumor microenvironment during breast cancer progression. While CK14 and CK5 consistently interact with each other representing the basal and luminal cell compartment, immune cells seem to increase their interaction with other immune cells (e.g., B Cells (CD20+)) and with cells potentially undergoing Epithelial-mesenchymal-transition (EMT) (e.g., Twist+ and Slug+)(Figure 5B). Next we plotted the first two components of the PCA of the results represented by their importance signatures to visualize how tumor grade (Figure 5C) and clinical subtypes (Figure 5D) are distributed.

**Figure 5.**
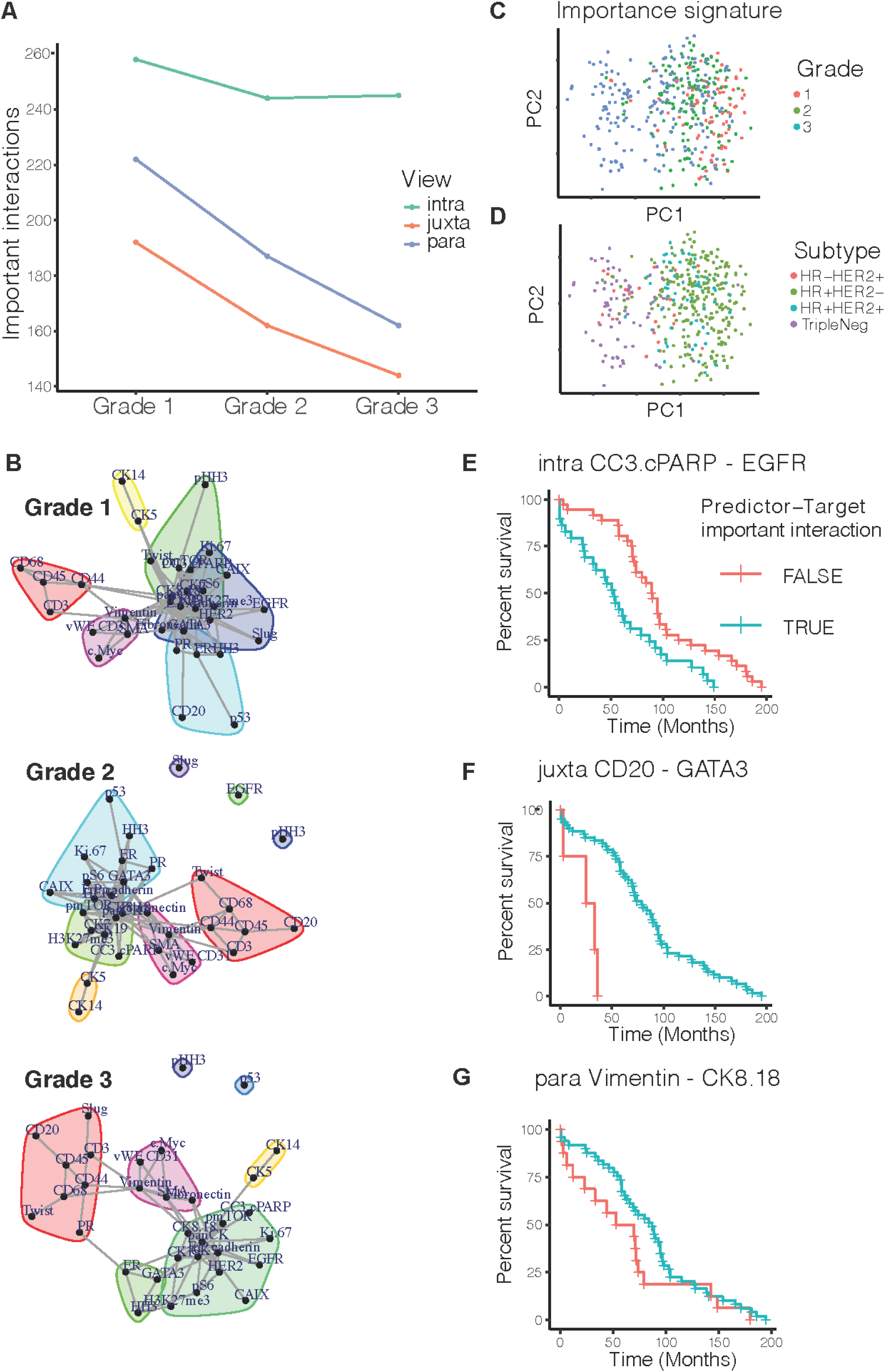
MISTy signatures can uncover clinically relevant features in IMC data from 415 breast cancer samples. (A) Change of total number of estimated important interactions per grade (Importance ≥ 0.5). (B) Changes in the tumor microenvironment can be visualized by network community plots representing the juxtaview for each tumor grade. E.g. the yellow cluster represents a constant link in the juxtaview between luminal-(CK8/18+) and basal-like (CK5+) cell types across all tumor grades, while the red cell cluster shows an increased interaction with tumor progression of immune cells (CD68+/CD45+), B cells (CD20+) and T cells (CD3+) with cells potentially undergoing EMT (Twist+/Slug+). (C) Importance signatures visualized as the first two components of a PCA highlight the separation of grade and (D) clinical subtype. Kaplan-Meier curves based on stratification by estimated importance of MISTy predictor-target interactions that were found to be correlated with patient outcome: (E) cleaved caspase 3 / cPARP and EGFR in the intraview; (F) CD20 and GATA3 in the juxtaview and (G) Vimentin and CK8/18 in the paraview.

We decided to focus our further analysis on grade 3 tumors only, since grade 1 and 2 samples are mostly annotated with HR+HER2-clinical subtypes (96% and 81% respectively) and the distribution of clinical subtypes for grade 3 samples is more balanced (47.6% HR−, 52.4% HR+ (out of 185), with the HR- group containing mostly triple negative subtype (79% out of 88 samples) and in the HR+ group 58,7% out of 97 samples are HR+HER2−). The importance signature for grade 3 clearly separated clinical subtypes (Sup.Figure 4A) based primarily on intraview for the first two components (Sup.Figure 4B), and we asked whether there are specific predictor-target interactions with high importance that could be linked to survival.

To associate MISTy results with clinical outcome, we calculated the Spearman rank correlation coefficients between the estimated importance of target-marker pairs to the overall survivability in months, selecting only pairs that contain at least 30% positive importance values. Samples from patients with multiple samples were treated as independent. Next, we performed the analysis accounting for clinical subtypes by running analysis on those samples independently. In the group of HR+HER2+ samples (n = 8), the estimated importance of 19 predictor-target pairs (5 intraview, 5 juxtaview, 9 paraview) is significantly correlated to the overall survival (p < 0.01). In the group of HR+HER2-samples (n = 20), the estimated importance of 25 predictor-target pairs (2 intraview, 8 juxtaview, 15 paraview) is significantly correlated to the overall survival and in the group of HR-HER2+ samples (n = 11), the estimated importance of 12 predictor-target pairs (3 intraview, 6 juxtaview, 3 paraview) is significantly correlated to the overall survival.

We focused specifically on the triple negative samples (n = 26), since currently no biomarkers are available that could be linked to outcome, where the estimated importance of 17 predictor-target pairs (9 intraview, 2 juxtaview, 6 paraview) is significantly correlated to the overall survival. We picked from the top predictor-target pairs correlated with overall survival for each view as an example for further analysis, but we provide all results for further experimental validation (Sup.Table 1). We stratified the samples by the estimated importance of the selected predictor-target interaction (cutoff 0.5). We then plotted the Kaplan-Meier curves and performed a log rank test to estimate the significance of the difference in overall survivability between the two groups. We found cleaved caspase 3 and cPARP, which are both markers of cell death, when estimated to interact (predictor-target) to EGFR in the intraview, is linked to better overall survival (*log*_10_*p* = − 2. 46, Figure 5E). In the juxtaview, we found that the absence of the interaction of B cells (CD20+) and cells of luminal phenotype (GATA+) are linked to bad overall survival (*log*_10_*p* = − 2. 86, Figure 5F). As a last example, we also found that estimated interactions between stromal cells (Vimentin+) and luminal luminal epithelial cells (CK8/18+) to be linked to better overall survival (*log*_10_*p* = − 2. 53, Figure 5G), which can be interpreted as an indication of tumor cell mass (less stroma - more tumor cells).

In summary, we could successfully link MISTy results and signatures to clinical features and survival outcomes. The provided list of features can be used as a resource for future experimental validations and, with an increasing amount of published spatial omics datasets linked to clinical data, we expect similar studies across various disease types and experimental technologies.

### Application to a spatial transcriptomics breast cancer dataset

Key features of MISTy are that it is technology agnostic and flexible to analyze different spatially-resolved data. Even more, the properties of the data obtained from different technologies can be leveraged to create different explanatory views.

To illustrate this, we analyzed the spatial gene expression profiles of two sections of a sample of invasive ductal carcinoma in breast tissue profiled with 10x Visium^36^. The 10x Visium slides contain 4992 total spots of 55 µm in diameter per captured area that enable the profiling of up to 10 cells per spot. With this technology, thousands of spatially resolved genes can be profiled simultaneously within a sample, allowing for characterization of molecular processes.

Previously, we have shown the utility of the footprint-based method PROGENy to robustly estimate the activity of signaling pathways, in both bulk and single-cell transcriptomics^37–39^. PROGENy estimates the pathways’ activity by looking at the expression changes in downstream target genes, rather than on the genes that constitute the pathway itself. Due to the resolution and the gene coverage of 10x Visium slides, the same approach can be applied to spatial transcriptomics datasets to enhance the functional view of the data. We estimated pathway activities for two reasons: 1) to reduce the dimensions of the data of each spot into interpretable and functionally relevant features, while still using the information of as many genes as possible, and 2) to provide a set of features that are more stable than the sparse expression of marker genes.

For each sample section, we estimated the activities of 14 cancer relevant signaling pathways of each spot using PROGENy^37,39^ (Figure 6A). While pathway crosstalk mechanisms are expected within a spot, we hypothesized that the local pathway activity could also be regulated by neighbouring cells in other spots to coordinate cellular processes. Therefore, we identified a set of 266 expressed genes in both sections annotated as ligands (Figure 6A) in the meta-resource OmniPath^40^ (see Methods) and designed a MISTy pipeline to model pathway activities using three different views: An intraview of pathway activities and two functional paraviews focusing on pathway activity and ligand expression, respectively (see Methods for full description of the views). Improvement in the prediction of pathway activities by this multi-view model would provide evidence of the relevance of intercellular communications in the regulation and maintenance of the functional state of a spot. Moreover, the traceable importances of each view may suggest possible mechanisms of intercellular communication.

**Figure 6.**
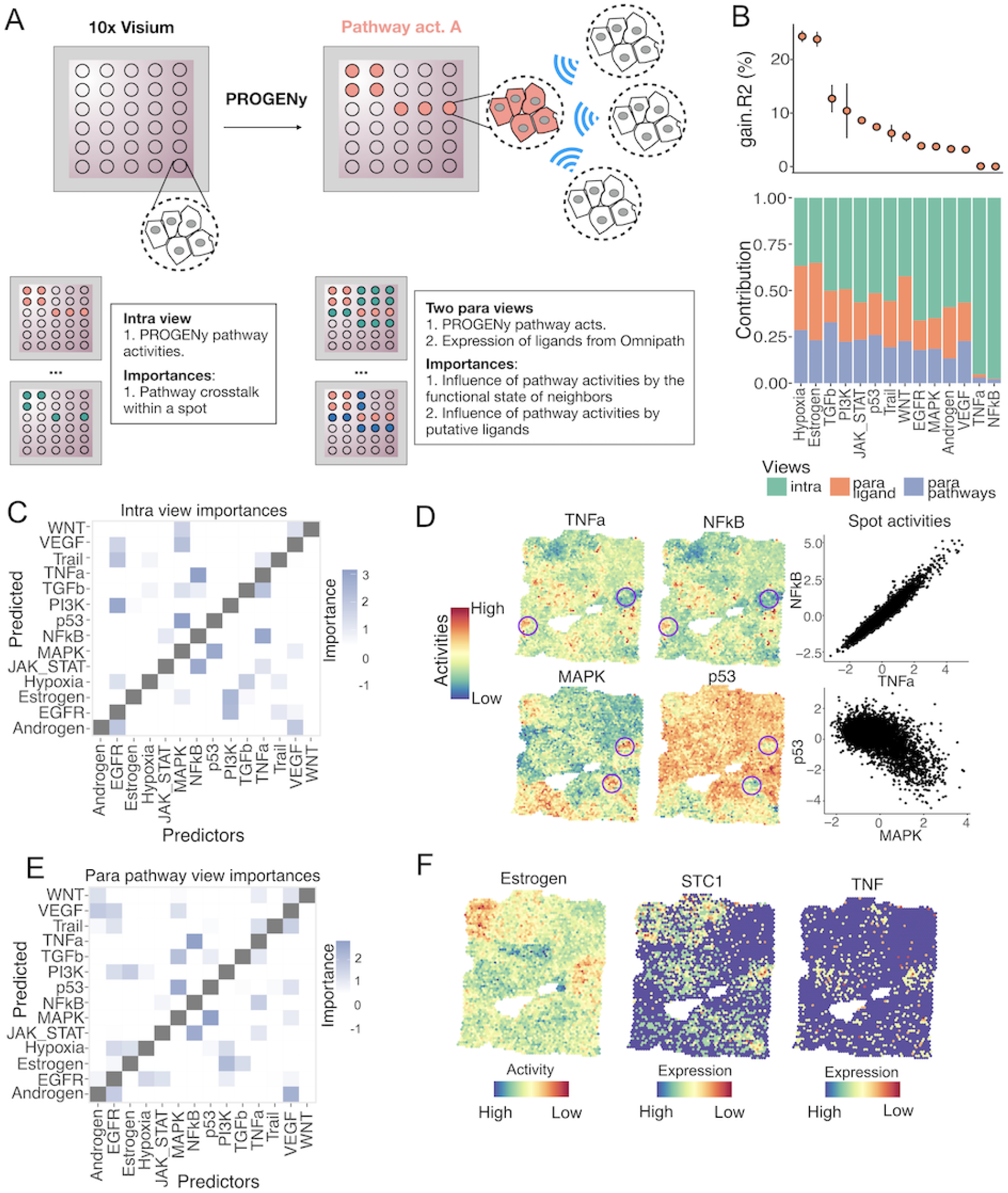
Application of MISTy to a spatial transcriptomics dataset. (A) Schematic of the MISTy pipeline used in Visium 10x slides. Each visium spot profiles the gene expression up to 10 cells. Pathway activities were estimated with PROGENy and a MISTy model was built to predict them using two spatially contextualized views. (B) Changes in R^2 observed in each predicted pathway after using the multiview model, reflecting the importance of the spatial context (upper panel). Contribution of each view to the prediction of the pathway activities in the meta-model. The stacked barplot represents normalized values of the fusion coefficients of the respective views for each pathway (lower panel). (C) Variable importances for the intraview. (D) Intrinsic associations of pathway activity scores of NFkB and TNFa, and p53 and MAPK. Spatial distribution of pathway activities from the first section. Circled areas exemplify niches where coordinated activities were observed. Scatterplots show the within-spot relationship between each pair of pathways. (E) Variable importances for the pathway paraview. (F) Spatial distribution of Estrogen pathway activities and scaled gene expression of SCT1 (top predictor of Estrogen in the ligand paraview) and TNF.

The multiview model improved significantly the variance explained of 11 of the 14 pathway activities (t-test on cross validations, mean adjusted p-value < 0.1), with improvements of up to 24% compared to the intraview model in the case of the estrogen and hypoxia pathways (Figure 4B). We found a mean contribution of 58% of the intraview, 22% of the ligand expression paraview and 20% of the pathway activity paraview to the prediction of pathway activities in the multi-view model (Figure 6B). We compared these results to the model performance in five iterations of slides with permuted layouts to provide further evidence of the importance of spatial information in the prediction of marker pathway activities (See Methods, Sup.Figure 6A,B). As expected, in these random slides we observed no improvements in variance explained when fitting models with spatially contextualized views (Sup.Figure 6A). Moreover, when we compared the view’s contributions of the models fitted to random and original slides, lower contributions of the paraviews were recovered for the random models (Sup.Figure 6B, Wilcoxon test *log*_10_*p* < − 10). These results confirmed that MISTy models are able to extract informative spatial relationships between markers in tissue samples where spatial organization is expected.

The importances of the features used as predictors in each view are consistent with biological processes. In the intraview (Figure 6C) we recovered, among others, associations between NFkB and TNFa, and P53 and MAPK that have been reported previously^37^. These results capture pathway crosstalks within a spot as illustrated in the spatial distribution of pathway activities shown in Figure 6D. Predictor importances in the pathway paraview captured similar associations as the ones captured by the pathway intraview (Figure 6E). The paraview importances, however, reflect patterns of tissue organization in which multiple neighboring spots share similar cellular states in larger areas. If a relationship between two pathways A and B is observed within a spot and a coordinated local activity of these pathways is happening, then the activity of pathway A of the neighbors of a given spot indirectly explains its pathway B activity. For example, the obtained paraview relationship between NFkB and TNFa, and P53 and MAPK (Figure 6E) explained the regions where a collection of spots showed coordinated higher or lower activities of these pathways (Figure 6D, circled areas). Additionally, new associations between pathways became relevant when taking into account the functional state of the neighbors of each spot (Figure 6E). In hypoxia, where the contribution of the para pathway view to the multi view model was 30%, estrogen and PI3K pathway activities had the highest importances, besides EGFR that was recovered from the intraview importances too (Sup.Figure 6C). The local expression of putative ligands contributed mostly to the prediction of estrogen, WNT and hypoxia (para ligand view contribution >= 34%, Figure 6B). We annotated each ligand-pathway interaction using Omnipath. We recovered the potential target receptors of all predictor ligands and assigned them to one of the 14 pathways in PROGENy based on the whole collection of annotations stored in Omnipath. Additionally, we annotated each predictor ligand as a direct by-product of a pathway if they belonged to one of the transcriptional footprints in PROGENy. From the 135 most important ligand-pathway interactions (importance >= 2), 84 could be annotated as described above. The 51 unannotated interactions could represent novel context dependent intercellular processes and show how MISTy could be used as a hypothesis generation tool. Among the top annotated interactions observed between the pathway activities and ligands (Sup.Figure 6D), we recovered the relationship between STC1 and estrogen pathway activities (Figure 6F). STC1 is a glycoprotein hormone that is secreted into the extracellular matrix and has been discussed in the literature as a promising molecular marker in breast cancer^41^. TNF, STC1’s reported receptor, showed similar spatial patterns in the slide (Figure 6F), suggesting a potential intercellular mechanism that mediates estrogen pathway activity. High importances to predict estrogen pathway activity were observed for other estrogen-receptor dependent genes such as EFNA1 and EDN1, as well as for the estrogen responsive gene TFF1 (Sup.Figure 6E). Interestingly, we observed that ligand importances clustered pathways that shared para pathway interactions, such as p53, MAPK, and TGFb. Altogether, our results showed that MISTy was able to improve the prediction of pathway activities by incorporating their spatial context. Moreover, we were able to identify known and novel spatial dependencies between pathway activities and ligands that reflect the functional organization of the tissue.

## Discussion

Here we present MISTy, an explainable framework for the analysis of highly multiplexed spatial data without the need of cell type annotation. It can scale and is technology agnostic enabling the analysis of increasingly complex data generated by recent and upcoming technologies. MISTy complements other methods that leverage spatial information to explore intercellular interactions. The current approaches focus mainly only on the local cellular niche, i.e., the expressions measured in the immediate neighborhood of each cell^26–29,31^.

Other methods that consider the broader tissue structure are relatively inflexible^30,32^. They consider a fixed form of nonlinear relationship between markers at predefined spatial contexts (e.g., fixed distance), and they do not scale well due to their high computational complexity. In contrast, MISTy offers a flexible range of spatial analyses in a scalable framework. We present a selected set of workflows for the analysis of spatial data, using not only the marker expressions but also derived features, such as pathway activities.

We established a performance baseline for MISTy on *in silico* data before applying MISTy to real-world data. We showed that MISTy achieves high performance on the task of reconstructing the intra- and inter-cellular networks of interactions.

We then applied MISTy to three real-world spatial-omics data sets from breast cancer samples. We applied MISTy on imaging mass cytometry data, capturing dozens of protein markers at (sub)cellular resolution. The results show that we were not only able to recapitulate results from the literature without prior-knowledge-based cell type annotation, but to also generate new hypotheses. Our results show that the information that is available from the expression of markers in the broader tissue structure is often more important than their expression in the local cellular niche. This highlights that not only cellular niches but also tissue structure have a direct impact on cellular states and should be included into the “microenvironment” definition. Furthermore, we show how MISTy finds interactions that are associated with clinical features.

Finally, we applied MISTy on a spatial transcriptomics data set measured with 10x Visium. Here, thousands of transcripts are measured in spots containing several cells. Given the richness of the data, we were able to go a step further and consider the analysis of functional features, in the form of pathway activities which were inferred from the data. In particular, we showed the crosstalk between pathways and the ligand-pathway interactions in the context of the broader tissue structure in breast cancer. Our results showed that MISTy in combination with functional transcriptomics tools and prior knowledge can be used in spatial transcriptomics to uncover coordinated functions that are maintained in niches of the tissue. Moreover, the explanatory component of the multiview model provides relevant predictors that could become the base of mechanistic models.

Although the interactions extracted by MISTy cannot be considered directly as causal, they can facilitate the downstream analysis of biological systems at the tissue level in several directions: (i) to predict the behavior of systems under perturbations, by using the MISTy model to generate marker expressions based on the new conditions; (ii) to guide the reconstruction of multicellular causal signaling networks, using databases to identify mechanisms giving rise to the extracted interactions; and subsequently (iii) to construct mechanistic models of the dynamical behavior of the system constrained by the extracted explanations.

The work we presented lays the foundation for further exploration of MISTy in several directions. One direction is to address the scalability of MISTy to millions of cells and thousands of markers per sample, which is beyond what the available technologies can offer, but is likely to come in the near future. To do this we are exploring approximate but accurate methods to replace the computationally expensive step of generating views where the pairwise distances between all cells need to be calculated. Another direction is the exploration of the performance that can be achieved by MISTy with different ensemble approaches using various types of explainable constituent models. Furthermore, MISTy can be used to generate more specific views. In particular, views that capture the spatial expression of specific cell types, so that we can dissect the spatial interactions between different cell types, or views that focus on regions of the tissue, for example, healthy vs pathological, where we would model the interactions between the functionally different regions. Finally, MISTy generates a model for each marker of interest that can be readily used to make predictions of marker expressions under different conditions. For example, we can increase or reduce the expression of a certain marker *in silico* and explore the effects of the new condition. Importantly, we can use the models to estimate how much the different views, such as intrinsic or paracrine effects, contribute to the prediction of the expression of each target marker.

In summary, we believe that MISTy is a valuable tool to analyse spatially resolved data, adaptable to multiple data modalities and biological contexts, that will also evolve as experimental techniques improve. An implementation of MISTy as a R package named mistyR (https://saezlab.github.io/mistyR/) is fully documented and freely available from GitHub, Bioconductor and as a Docker image.

## Methods

### *In silico* benchmark

The *in silico* model is a two-dimensional cellular automata model that focuses on signaling events, therefore cell growth, division, motility and death are neglected. First, we created ten random layouts. To account for cellular heterogeneity in the tissue, we assigned one of four different cell-types *CT*1, ... *CT*4, to each spot of the layout or left it empty (intercellular space). Each of these cell-types has a distinct set of receptors expressed and distinct intracellular wiring. To keep the model simple, we considered 11 biological species S = {ligands: ligA, ligB, ligC and ligD; intracellular proteins: protE, protF; ligand producing nodes: prodA, prodB, prodC and prodD}. The intracellular processes involve two layers, first the ligand activation of signaling hubs, represented by protE and protF, and ligand production/secretion regulated by proteins (prodA, prodB, probC and prodD)(Figure 2A). The model simulates the production, diffusion, degradation and interactions of these 11 molecular species on a 100-by-100 grid. Ligands are produced in each cell-type based on the activity level of their production nodes and then freely diffuse, degrade or interact with other cells on the grid. Other molecular species involved in signaling are localised in the intracellular space and their activity depends on ligand binding and intracellular wiring.

The model is formally stated by the following partial differential equations for each species:

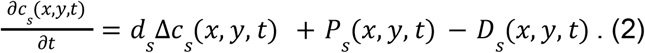

This equation describes the diffusion, the production/activation and the degradation of the species. We made the following assumptions. *c*_*S*_(*x*, *y*, *t*) is the concentration of species *s* ∈ *S* at the grid point (x,y) at time t. The diffusion is homogenous across the image, the diffusion coefficient of species *s* is *d*_*S*_. Only ligands and ECM are diffusing, other intracellular molecules cannot leave the cell.

The production term includes the generation of ligands and the activation of intracellular proteins. It depends on the cell-types and the activity of the production node: for ligands, *ligX* production depends on *prodX* node: *P*_*i*_(*x, y, t*) = α_*i,ct*_*prodX*_*i*_(*x, y, t*) for *i* ∈ {*ligA*, *ligB*, *ligC*, *ligD*} and *ct* ∈ *CT*, the α_*i,ct*_ coefficient defines which cell type produces which proteins and how strongly the production depends on the activity of the production node.

For intracellular proteins, the protein activity depends on the abundance of the ligands that activate its pathway: 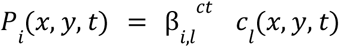, for *i* ∈ {*protE*, *protF*}and *l* ∈ {*ligA*, *ligB*, *ligC*, *ligD*}. The 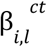 encodes the interactions between the ligand and the proteins in each cell type..

Degradation is proportional to the concentration of ligands, intracellular proteins and ECM, *D*_*S*_(*x, y, t*) = *γ*_*S*_*C*_*S*_(*x, y, t*), where *γ*_*S*_ is a constant degradation coefficient.

The above model was simulated from a randomised initial condition until a steady state distribution of the species and activity was achieved (Sup.Figure 1). We considered the steady state values of {ECM, prodA, prodB, prodC, prodD, protE and protF} as available measurements for the MISTy workflow. In contrast to the included measurements, our assumption is that the expression of ligands would be more difficult to capture in a real experiment and therefore excluded them.

We aggregated the interactions from the mechanistic model for the different cell types in joint binary matrices of directed ground truth interactions for the different views. To compare the matrices to the importance matrices from the output of MISTy, we transformed the joint matrices into undirected matrices *A*_*u*_ = *sgn*(*A* + *A*^*T*^). We then quantified the performance of MISTy for the task of reconstruction of intra- and intercellular networks from the true and the extracted interaction matrices.

### Data acquisition and processing

#### Imaging Mass Cytometry

The first Imaging Mass Cytometry dataset consists of 46 samples from 26 breast cancer patients with varying disease grades^28^. The original data consisted of 50 samples, from which we removed samples coming from normal tissue. The raw data was segmented and single cell features were extracted with histoCAT. The samples contain between 267 and 1455 cells with measured expression of 26 proteins / protein modifications. The cell-level data was preprocessed as defined in Arnol et al.^32^ in order to assure the validity of direct comparison of results.

The second Imaging Mass Cytometry dataset consists of 720 samples from 352 breast cancer patients from two cohorts, with long term survival data available for 281 of those patients^35^. The samples contain measurement of 37 proteins / protein modifications. The raw data was segmented and single cell features were extracted with histoCAT. The cell-level data was preprocessed as in the original study. To ensure robustness of the results, we filtered samples containing less than 1000 cells, samples coming from a normal or control tissue and samples without annotated tumor grade or clinical subtype, resulting in a total of 415 samples for our analysis.

#### Spatial transcriptomics

The data and sample information were obtained from 10x Genomics^36^. The data consists of spatial transcriptomic measurements of two sections of a sample analyzed with 10x Genomics Visium. The sections come from tissue from a patient with grade 2 ER^+^, PR^−^, HER2^+^, annotated with ductal carcinoma in-situ, lobular carcinoma in-situ and invasive carcinoma. The mean sequencing depths were reported to be 149,800 and 137,262 reads per spot for a total of 3,813 and 4,015 spots per section respectively. The median UMI counts per spot were reported as 17,531 and 16,501, and the median genes per spot as 5,394 and 5,100 respectively. The raw data was preprocessed and count matrices were generated with *spaceranger-1.0.0*. Individual count matrices were normalized with *sctransform* implemented in Seurat 3.1.2^42^. For each spot, we estimated signaling pathway activities with PROGENy’s model matrix using the top 1000 genes of each transcriptional footprint. We retrieved from Omnipath^40^ all proteins labeled as ligands and in each dataset we filtered all ligands whose expression was captured in at least 5% of the spots.

### View generation

For the application of MISTy on IMC and spatial transcriptomics data, in addition to the intraview, we considered creating views by aggregating the available spatial and expression information in two ways. For the IMC data we created a view that describes the local cellular niche (juxtaview) and a view that describes the broader tissue structure (paraview), both created using the measured marker expression directly. For the spatial transcriptomics data we created a paraview using the estimated pathway activities at each patch and a paraview using the measured expressions of a set of ligands.

We consider a dataset *D* = [*X*_*(n×s)*_ *Y*_*(n×k)*_], represented as a matrix of dimensions *n* × (*s + k*) of spatially-resolved highly-multiplexed measurements of a sample, where *n* is the number of measured units (pixels, cells, patches) available in the sample, *s* is the number of spatial dimensions in the geometry matrix *X* and *k* is the number of measured markers in the expression matrix *Y*.

The juxtaview was generated by summing the expressions of its direct neighboring cells, i.e., 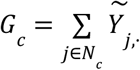, where *N*_*c*_ represents the set of neighboring cells of cell *c*. The neighboring cells for each cell can be determined either during image segmentation, for example by setting a threshold of membrane to membrane distance, or, as in the case for the application of MISTy on IMC data, by post-hoc neighborhood estimation. For the application of MISTy on IMC data the neighborhood of each cell in a sample was estimated by constructing a cell graph by 2D Delaunay triangulation followed by removal of edges with length larger than the 25th percentile of all pairwise cell distances across all samples, which corresponded to 11.236 microns from the cell centroid.

The paraview was generated by weighted aggregation of the expressions of all cells (patches) from the sample 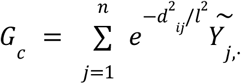, where *d*_*ij*_is the Euclidean distance between cells *i* and *j*, calculated from matrix *X* and *l* is a parameter controlling the shape of the weighting radial basis function. The parameter *l* corresponds to the radius around the cell where we consider the values of the weighting function to allow for substantial contribution of the measured expressions. For the application of MISTy to both IMC and spatial transcriptomics data, we optimized the value for the parameter *l*. For each IMC sample, we constructed models for each marker, with parameter *l* ∈ {25, 50, 100, 200, 400}. This corresponds to an effective radius of influence of 25 to 400 pixels or micrometers. The mean values of the parameter l across all samples for all markers is shown in Sup.Figure 5. Given the resolution of 10x Visium, MISTy models for spatial transcriptomics were built for each pathway activity using as paraview parameter *l* ∈ {2, 5, 10}, corresponding to a radius of influence of up to 10 spots. Then, for each marker we selected the value for *l* such that the estimated improvement in predictive performance by using the multi-view model in contrast to the intraview model is maximized. For each model (Random Forest), we estimate the predictive performance by measuring the variance explained on out-of-bag samples.

### Importance weighting and result aggregation

To calculate the interaction importances from a sample we used information from the two layers of interpretability and explainability: the values of the fusion parameters (α_*v*_ in Eq. (1)) in the meta-model and their respective p-values *p*_*k*_^*(v)*^ for each target marker *k* and the importances *I*_*kj*_^*(v)*^ of features *j* for the prediction of each target marker extracted from the predictive model for view *v* yields the MISTy interaction importance:

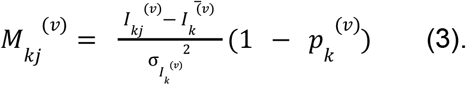

Since the importances *I*_*kj*_^*(v)*^ extracted from a Random Forest model (used for the current instance of MISTy) represent the amount of variance reduction in the target expression, the MISTy interaction importances correspond to the standardized value (by mean 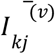 and variance 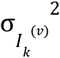) of the variance reduction weighted by the quantile 1 − *p*_*k*_^(*v*)^ of the statistic under the null hypothesis of zero contribution of the fusion coefficient for view *v* for target *k* in the linear meta-model. The importance derived from variance reduction can be generalized to any measure of impurity or values extracted by other feature importance estimation methods, given the model constituents of MISTy. Since the MISTy importances are standardized, importances from multiple samples can then be aggregated by simple averaging, while their interpretation remains the same.

For views that contain the same set of predictors as targets, we also identified the communities of interactions from the estimated importances. For this, we transformed the square matrix *A* of estimated predictor-target interactions to an undirected graph adjacency matrix as *A*_*p*_ = *A* + *A*^*T*^. We then extract the community structure from the graph using the Louvain algorithm^43^, a commonly used algorithm for community detection by grouping nodes, such that the modularity of the graph is maximized.

### Permutation of the slides

To evaluate the performance of MISTy models in samples with no spatial organization, we generated random samples for both the IMC and spatial transcriptomics data and ran the same pipeline as for the original data. We permuted the coordinates of each spatial unit (cell or spot) for each slide ten times for the IMC data and five times for the spatial transcriptomics data. Results were grouped and compared to the ones obtained in the slides with the original spot layout.

### Annotation of predictive ligands from the spatial transcriptomics pipeline

To facilitate the interpretation of the paraview ligand importances observed in the spatial transcriptomics models we assigned each ligand to a PROGENy pathway in two ways: 1) as a byproduct of pathway activation or 2) as a potential activator of a pathway. A ligand was considered as a byproduct of a PROGENy pathway if it was part of its top 1000 footprint genes. A ligand was considered as a putative activator of the pathway it predicted if at least one of its possible receptors could be assigned to it. We used OmnipathR to annotate each receptor using the *import_omnipath_annotations* function and regular expressions to filter annotations associated to the PROGENy pathway of interest. Only ligands with a paraview importance >=2 were considered in this annotation (Sup.Figure 6D).

## Acknowledgements

J.T. acknowledges the financial support from the European Union and the Slovenian Ministry of Education, Science and Sport (agreement No. C3330-17-529021). D.S. was supported by an Early Postdoc Mobility fellowship (no. P2ZHP3_181475) and was a Damon Runyon Fellow supported by the Damon Runyon Cancer Research Foundation (DRQ-03-20). We would like to thank Nicolàs Palacio-Escat for the Figure 1 design and Olga Ivanova for providing feedback on the manuscript.

## Author contributions

Conceptualization - J.T. and J.S.R.; Formal analysis - J.T.; Investigation - J.T. and D.S; Methodology - J.T.; Software - J.T., A.G. and R.O.R.F.; Supervision - J.S.R.; Visualization - J.T., R.O.R.F and A.G.; Writing - original draft - J.T., A.G and R.O.R.F; Writing - review & editing - D.S., and J.S.R.

## Supplementary material

**Sup.Figure 1.**
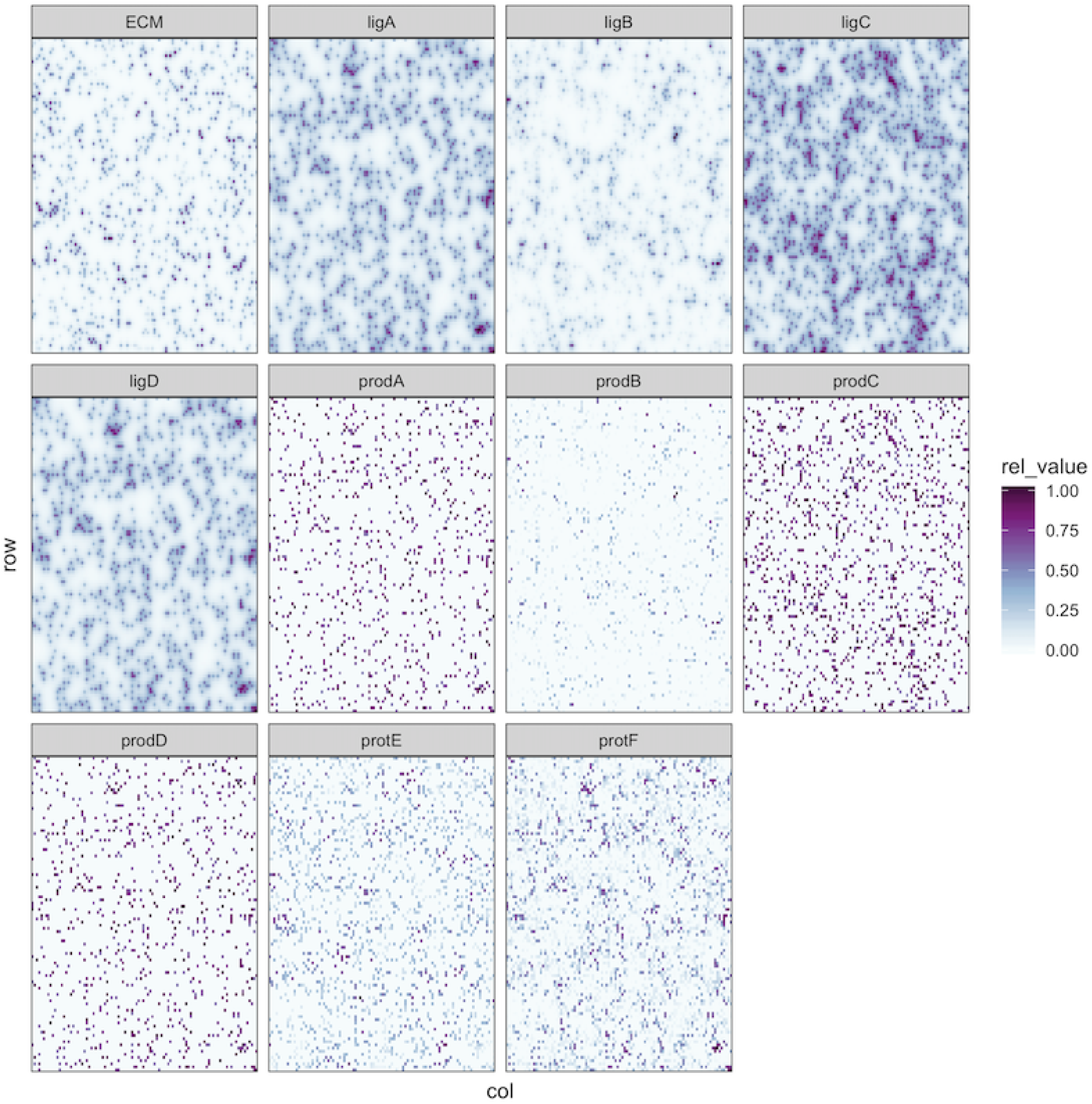
Steady state distribution of simulated species from a single sample.

**Sup.Figure 2.**
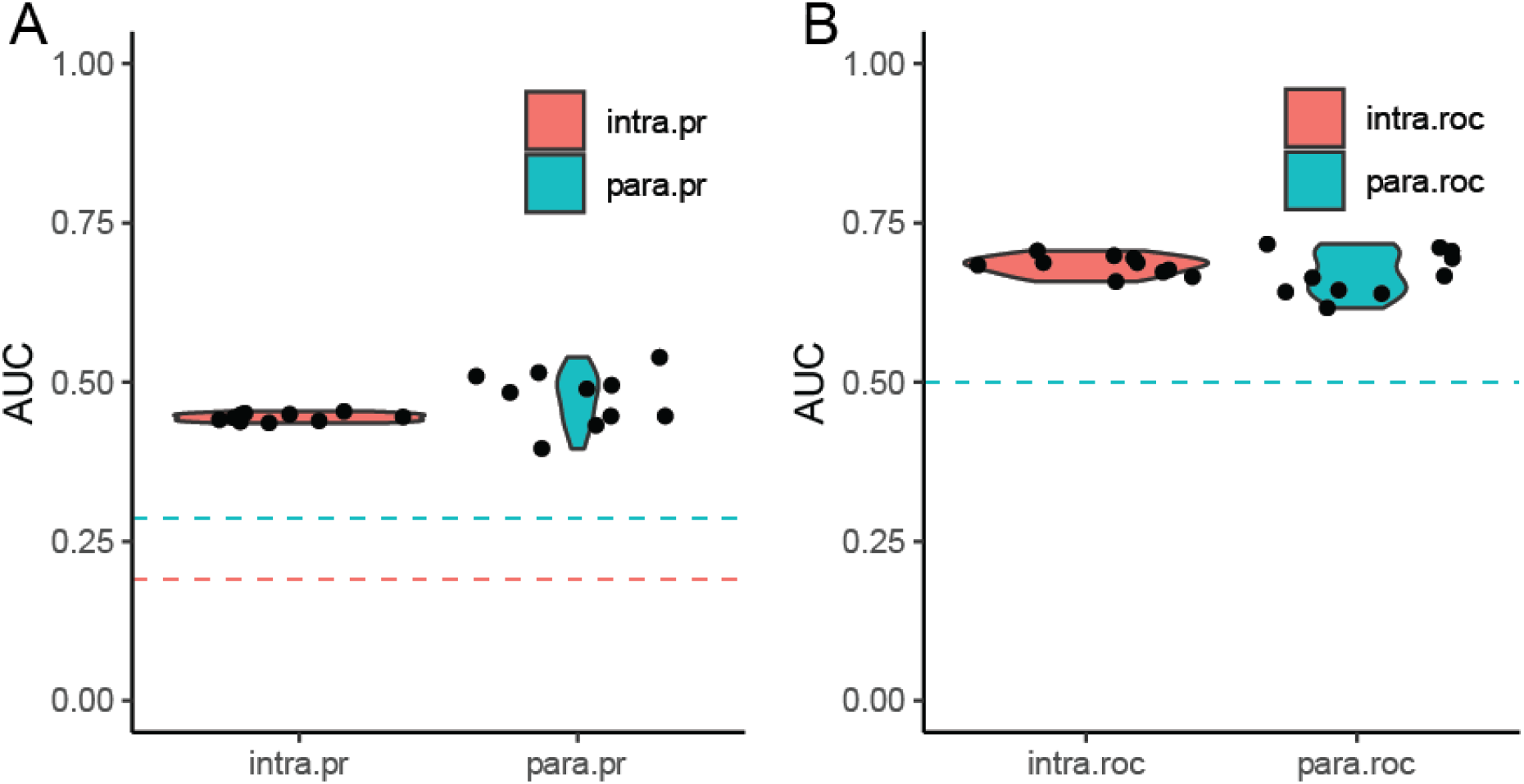
Distribution of Area Under the Precision-Recall curves (AUPRC, panel A) and Receiver-Operating Characteristic curves (AUROC, panel B) for the in-silico case study.

**Sup.Figure 3.**
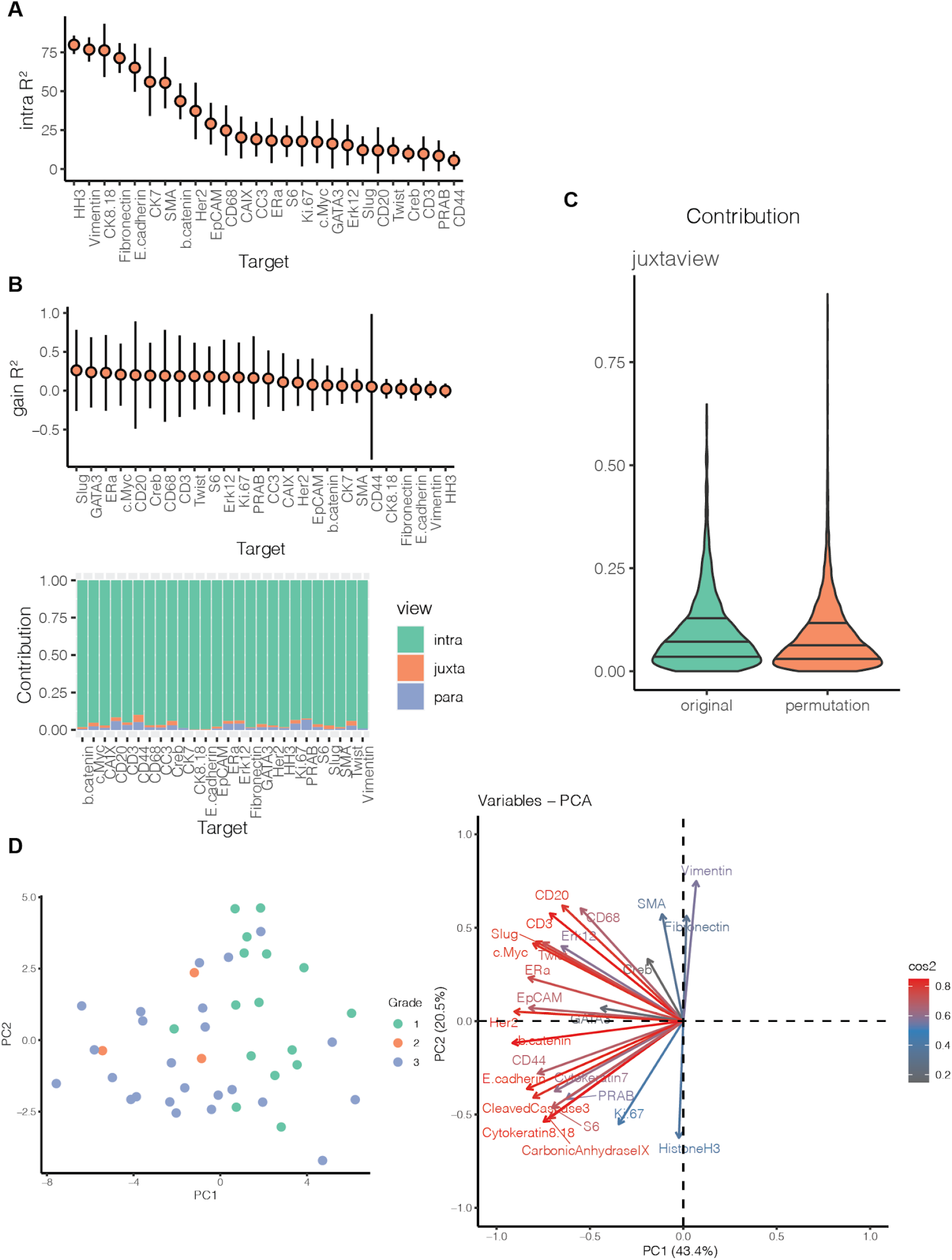
Additional plots for the IMC data from 46 breast cancer samples. (A) Predictive performance (variance explained) for all samples when considering intraview only (in absolute percentage points). (B) Gain in variance explained and relative contribution of each view to the prediction of the expression of the markers for samples with 10 random permutations of the cell locations. (C) Distribution of the relative contribution of the juxtaview to the prediction of the markers across all markers and samples with original cell locations and 10 random permutations. (D) First two principal components of the samples represented by the mean expression of the markers across all cells and the importance of the mean expression of the markers in the principal component analysis.

**Sup.Figure 4.**
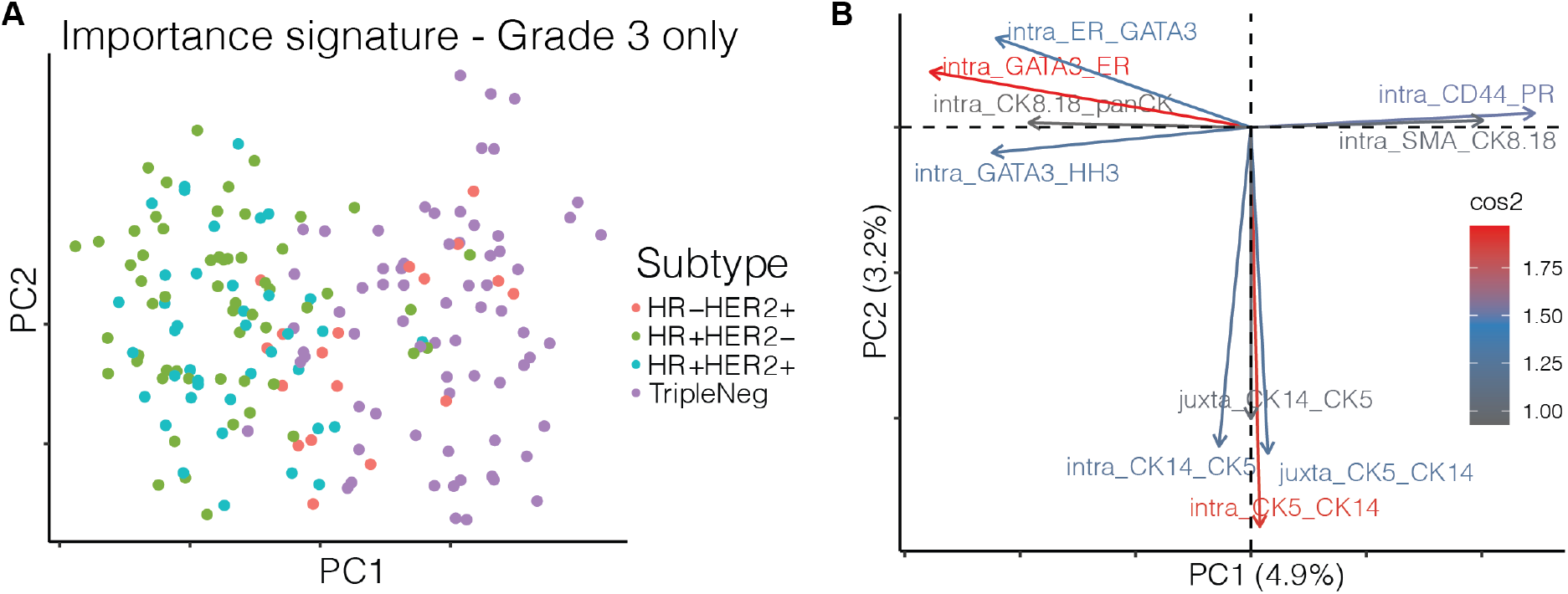
Importance signature and contrasts of IMC data from 415 breast cancer samples. (A) First two principal components of the importance signature of the samples and (B) importance of the variables of the signature in the principal component analysis (10 variables with the highest square cosine shown).

**Sup.Figure 5.**
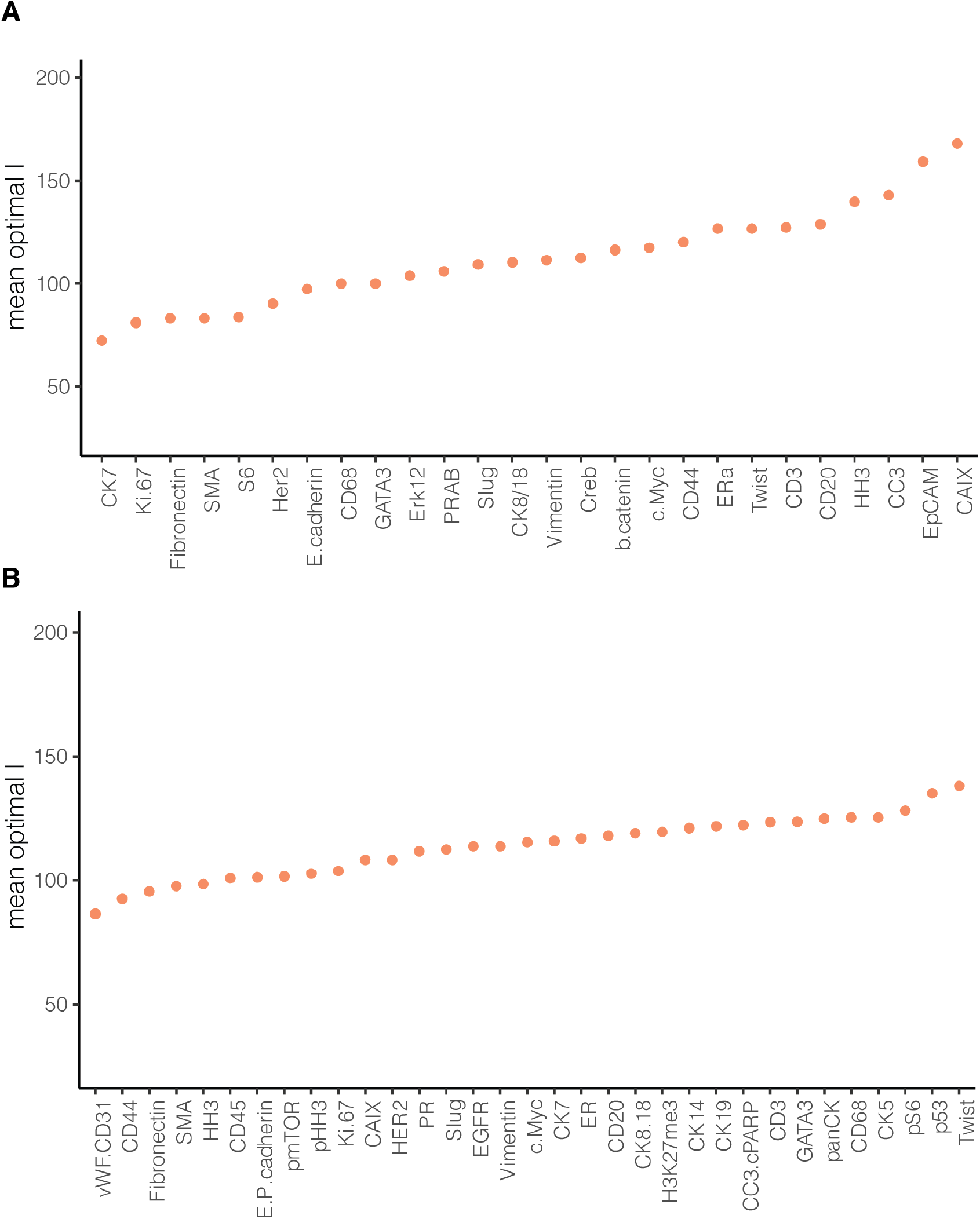
Estimated optimal value for the parameter l. (A) for the IMC data from 46 breast cancer samples and (B) for the IMC data from 415 breast cancer samples.

**Sup.Table 1.** Correlation of estimated importances to overall survivability for triple negative samples from the IMC data with 415 breast cancer samples.

| Predictor | Target | Spearman $\rho$ | $\log_{10} p$ | View |
| --- | --- | --- | --- | --- |
| H3K27me3 | Twist | -0.801 | -5.04 | intra |
| CC3.cPARP | EGFR | 0.605 | -2.46 | intra |
| Twist | HH3 | -0.595 | -2.38 | intra |
| Vimentin | HH3 | -0.588 | -2.33 | intra |
| CC3.cPARP | CAIX | 0.586 | -2.31 | intra |
| CC3.cPARP | Slug | 0.574 | -2.22 | intra |
| CD44 | GATA3 | 0.574 | -2.22 | intra |
| CK7 | panCK | -0.566 | -2.16 | intra |
| H3K27me3 | CK19 | -0.563 | -2.14 | intra |
| CC3.cPARP | Twist | 0.557 | -2.10 | intra |
| pS6 | EGFR | 0.555 | -2.08 | intra |
| Vimentin | Ki.67 | -0.666 | -3.01 | juxta |
| CD20 | GATA3 | -0.650 | -2.86 | juxta |
| Fibronectin | Slug | 0.613 | -2.53 | juxta |
| E.P.cadherin | pHH3 | 0.597 | -2.40 | juxta |
| CK8.18 | CC3.cPARP | -0.569 | -2.18 | juxta |
| HER2 | Fibronectin | 0.549 | -2.04 | juxta |
| pS6 | CD68 | -0.667 | -3.02 | para |
| Vimentin | CK8.18 | -0.613 | -2.53 | para |
| CD45 | p53 | 0.593 | -2.37 | para |
| E.P.cadherin | Twist | 0.566 | -2.16 | para |

**Sup.Figure 6.**
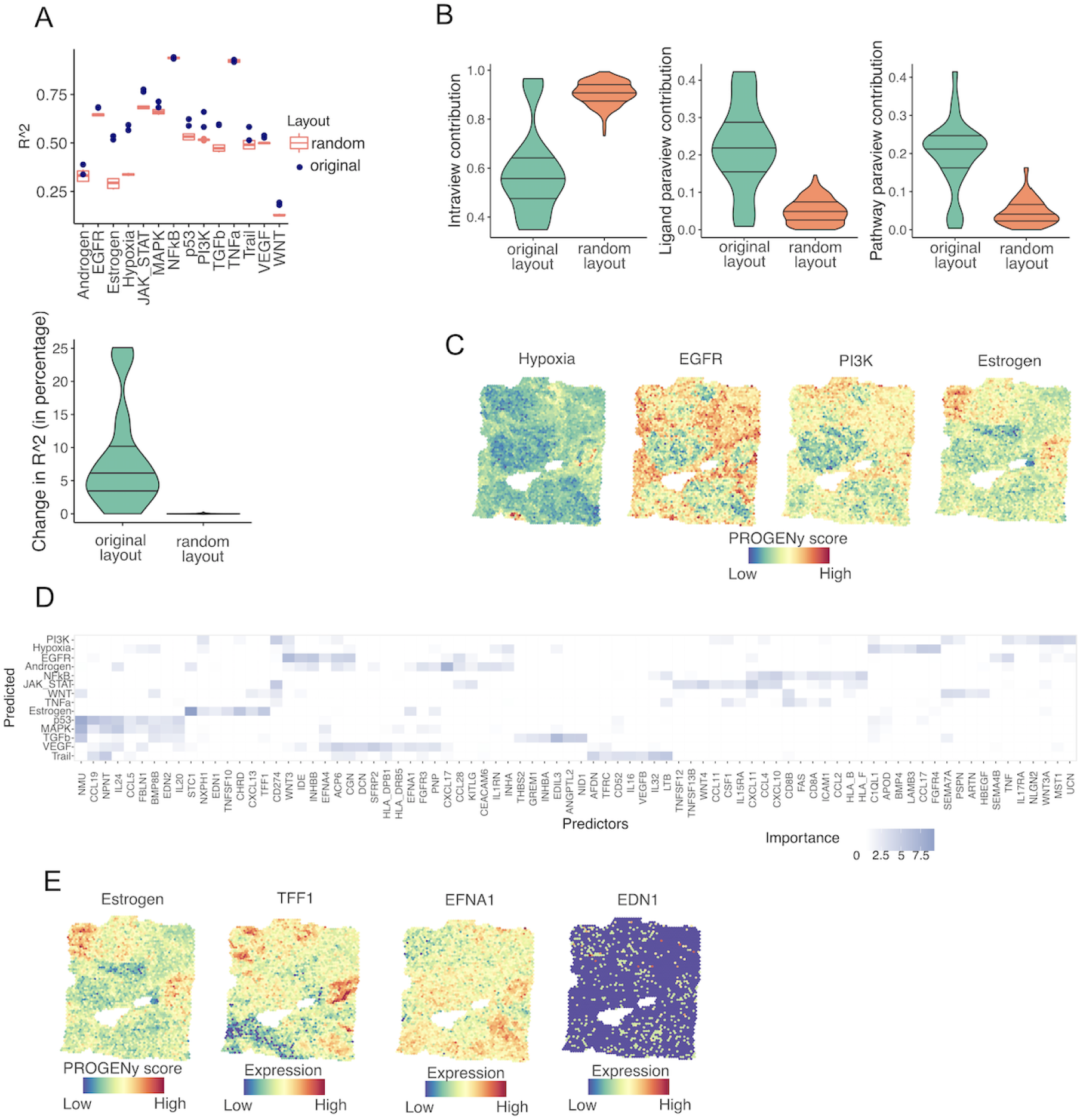
Application of MISTy to a spatial transcriptomics dataset. (A) Comparison of MISTy’s performance between original slides and a collection of 10 random slides with permuted coordinates. Differences of the R^2 of the multiview model (upper panel) and changes in R^2 between single and multiview models of both classes of models; (B) Differences in view contributions of all predicted markers between models fitted to original or permuted data. (C) Spatial distribution of pathway activities of Hypoxia and its top predictors in the paraview: EGFR, PI3K and Estrogen (D) Variable importances for the ligand paraview. (E) Spatial distribution of Estrogen pathway activities and scaled gene expression of TFF1, EFNA1, and EDN1 (additional top predictors in the ligand paraview).

## Notes

### Competing Interest Statement

The authors have declared no competing interest.

